# Metal-Induced Energy Transfer (MIET) for Live-Cell Imaging with Fluorescent Proteins

**DOI:** 10.1101/2022.11.12.516247

**Authors:** Lara Hauke, Sebastian Isbaner, Arindam Ghosh, Isabella Guido, Laura Turco, Alexey I. Chizhik, Ingo Gregor, Narain Karedla, Florian Rehfeldt, Jörg Enderlein

**Author notes:** Corresponding authors: N.K. Narain Karedla; F.R.; J.E. Equal contribution first authors (Lara Hauke, Sebastian Isbaner, Arindam Ghosh).

## Abstract

Metal-Induced Energy Transfer (MIET) imaging is an easy-to-implement super-resolution modality that achieves nanometer resolution along the optical axis of a microscope. Although its capability in numerous biological and biophysical studies has been demonstrated, its implementation for live-cell imaging with fluorescent proteins is still lacking. Here, we present its applicability and capabilities for live-cell imaging with fluorescent proteins in diverse cell types (adult human stem cells, human osteo-sarcoma cells, and *Dictyostelium discoideum* cells), and with various fluorescent proteins (GFP, mScarlet, RFP, YPet). We show that MIET imaging achieves nanometer axial mapping of living cellular and sub-cellular components across multiple timescales, from a few milliseconds to hours, with negligible phototoxic effects.

## 1 Introduction

Fluorescence nanoscopy^1^ beyond the classical diffraction limit of optical microscopy has become an indispensable tool for modern life sciences, allowing to discern the spatial organization of biological structures down to molecular length scales. The first successful method of super-resolution microscopy was stimulated emission depletion (STED) microscopy,^2^ which was followed by the big family of single-molecule localization microscopy (SMLM) techniques. The latter comprises photo-activated localization microscopy (PALM),^3^ stochastic optical reconstruction microscopy (STORM),^4^ fluorescent PALM (fPALM),^5^ direct STORM (dSTORM),^6^ and point accumulation for imaging in nanoscale topography (PAINT)^7^. These methods routinely achieve lateral resolutions down to a few dozen nanometers.

Achieving a comparable resolution along the third dimension (the optical axis) requires additional modifications of these methods: For STED, one has to apply special phase plates to generate an optical bottle along the optical axis,^8^ and for SMLM, methods such as bi-plane imaging,^9^ astigmatic imaging,^10^ wavefront shaping,^11^ or single-molecule self-interference^12^ have been developed for three-dimensional single molecule localization. Remarkably, the axial resolution achieved by all these methods is typically by a factor of 3-5 worse than the lateral resolution, very similar to the situation encountered in classical diffraction-limited confocal microscopy.

To overcome this anisotropic resolution, interferometric methods such as iPALM^13^ and isoSTED^14^ have been developed. These methods use two opposing objective lenses to image the sample, similar to what is done in diffraction-limited 4*π* microscopy^15^. While achieving truly isotropic super-resolution down to a few nanometers, these methods are technically highly complex which prevented their wide application so far. This is also true for the latest addition to the pool of super-resolution methods, MINFLUX^16^, which uses a complex three-dimensional triangulation method for pinpointing the position of a single molecule with unprecedented and even isotropic resolution^17^.

For achieving axial super-resolution close to an interface (∼200-300 nm axial distance), a different family of super-resolution methods has been developed that rely on electromagnetic near-field effects. The first one is variable-angle total internal reflection fluorescence (vaTIRF) microscopy^18^. It is based on conventional TIRF microscopy, but records several consecutive images of the same sample under different incidence angles of the totally reflected excitation light. This incidence angle variation generates evanescent excitation intensities with varying decay length. By applying an appropriate mathematical analysis, the axial distance of fluorescence structures from the glass surface can be recovered^19^. The second of these near-field methods is supercritical angle fluorescence microscopy (SAF microscopy)^20^, which uses the fact that the electromagnetic near-field of an emitting fluorescent molecule, which usually does not take part in its far-field emission (i.e. which does not contribute to observable fluorescence intensity), can couple into propagating and detectable far-field modes in the coverslide glass when the emitter comes sufficiently close to the glass surface (few hundred nanometers). These near-field-generated propagating modes travel at angles above the critical angle of total internal reflection and are thus called super-critical fluorescence (SAF). In contrast, the usual far-field emission of the molecule does strictly travel at angles below this critical angle (undercritical angle fluorescence or UAF). While the SAF emission intensity is strongly dependent on the distance of a molecule to the glass surface, the UAF emission intensity is independent of it. By measuring the ratio of both intensities, the axial position of an emitter can be determined with nanometer accuracy^21–23^.

Metal-induced energy transfer (MIET) imaging exploits similar physics for achieving axial super-resolution^24^. It uses the coupling of the near-field modes of a fluorescent molecule into so-called surface plasmons (collective metal electron oscillations) in a thin metal layer (typically 20 nm gold film) deposited on top of a coverslip surface. This leads to a strong distance-dependent modulation of the fluorescence lifetime of the emitter which can be used to convert its measured lifetime into an axial position^25^. Because MIET is based on lifetime measurements, it is more robust against intensity-affecting artifacts that may impact vaTIRF or SAF microscopy results. Moreover, provided the availability of a fluorescence lifetime imaging microscope (FLIM), MIET does not require any modifications of the microscope setup except that it needs glass coverslips covered with a thin metal layer, which can be routinely produced today with chemical vapor deposition. Similar to vaTIRF and SAF microscopy, MIET has an operating distance range of ∼150 nm - 200 nm from the metal surface where it can achieve an axial resolution down to a few nanometers. Previous applications of MIET imaging include basal membrane mapping in living cells^24^, three-dimensional reconstruction of focal adhesions and stress fibers^26^, the nuclear envelope architecture^27^, visualizing the dynamics of epithelial-mesenchymal transitions (EMT)^28^, single-molecule localization and co-localization^29, 30^, mapping the basal membrane and lamellipodia of human aortic endothelial cells^31, 32^, and, in combination with SMLM, three-dimensional isotropic resolution imaging of microtubules and clathrin pits^33^. Recently, it was shown that substituting the metal layer with a single sheet of graphene (graphene-induced energy transfer or GIET imaging) achieves a ca. tenfold higher axial resolution than MIET (down to a few Ångströms), within a distance range of ∼25 nm from the graphene^34–37^.

So far, all MIET/GIET applications mentioned above were done exclusively with synthetic organic dyes as fluorescent labels. For applying MIET to live-cell super-resolution microscopy it is important that MIET works with fluorescent proteins in living cells. Fluorescent proteins are widely used to label cellular structures of interest and are conveniently expressed within genetically modified living cells^38^. However, applying MIET to live cell imaging with fluorescent proteins is much more challenging than using MIET with synthetic organic dyes. Firstly, one has to assure that the average fluorescence lifetime values of the used proteins is homogeneous throughout the cellular structures (in the absence of any quenching metal layer). Secondly, one has to cope with the usually non-mono-exponential fluorescence decay of fluorescent proteins^39^. In the present paper, we demonstrate that fluorescent proteins (in particular, wild-type GFP and mScarlet) provide indeed fluorescent labeling with sufficiently homogeneous lifetime distributions across living cells, which is a necessary prerequisite for MIET, and that the error introduced by the non-mono-exponential decay of the fluorescent proteins into the lifetime-to-distance conversion still allows for an axial resolution of approximately 10 nm. We demonstrate fluorescent-protein MIET imaging of actin stress fibers in living mammalian cells, and of the basal membrane dynamics of *Dictyostelium discoideum (D*.*d*.) cells with nearly video-rate of imaging. Our work demonstrates the suitability of MIET for superresolution microscopy of living cells and sub-cellular components labeled with fluorescent proteins while keeping photo-toxicity to a minimum and achieving high temporal resolution. This is of paramount importance for the application of MIET to a wide range of biologically important problems.

## 2 Results

Fluorescence intensity and lifetime images were recorded with a home-built confocal laser-scanning microscope, see Figure 1A and section *Methods* for details. This microscope allows for taking live-cell fluorescence lifetime images with video-rate acquisition speed (fluorescence lifetime imaging microscopy or FLIM). The first system that we studied was live cells of the slime mold *D*.*d*., a social amoeba growing in soil and a model organism for investigating cell adhesion, motility, chemotaxis, and signal transduction^40^. Binding of cyclic AMP (cAMP) to the cells’ cyclic AMP receptor 1 (cAR1) results in amplification of actin polymerization at the leading edge of *D*.*d* cells, which leads to the formation of membrane protrusions known as pseudopodia^41, 42^. The cell line used in this work was derived from the axenically growing strain *D*.*d. carA-GFP*. MIET imaging was done for cells at the vegetative and development stages (see ‘Methods’ for detailed description of cell culture and transfection protocols). Unlike mammalian cells, *D*.*d*. cells lack integrin. Since they can bind equally well to hydrophobic and hydrophilic surfaces, adhesion is likely governed by van der Waals interactions of membrane glycoproteins with the substrate ^43^. Additionally, cell-substrate adhesion changes during the developmental time of *D*.*d*. cells ^44^. The *D*.*d*. cells in their developed stage that were used in this study were starved for 6 hours and pulsed with 50 nM cAMP every 6 minutes over the duration of starvation. The pulsing renders the cells chemotactic, which makes them polarized and they can migrate faster towards the source of cAMP. As a result, these cells show rapid motion and membrane dynamics which makes them ideal systems for checking the live-cell imaging capabilities MIET. Panels B and C in Figure 1 show fluorescence intensity and corresponding lifetime images of a representative *D*.*d*. cell where cAR1 in the cell membrane were labeled with green fluorescent protein (GFP).

**Figure 1:**
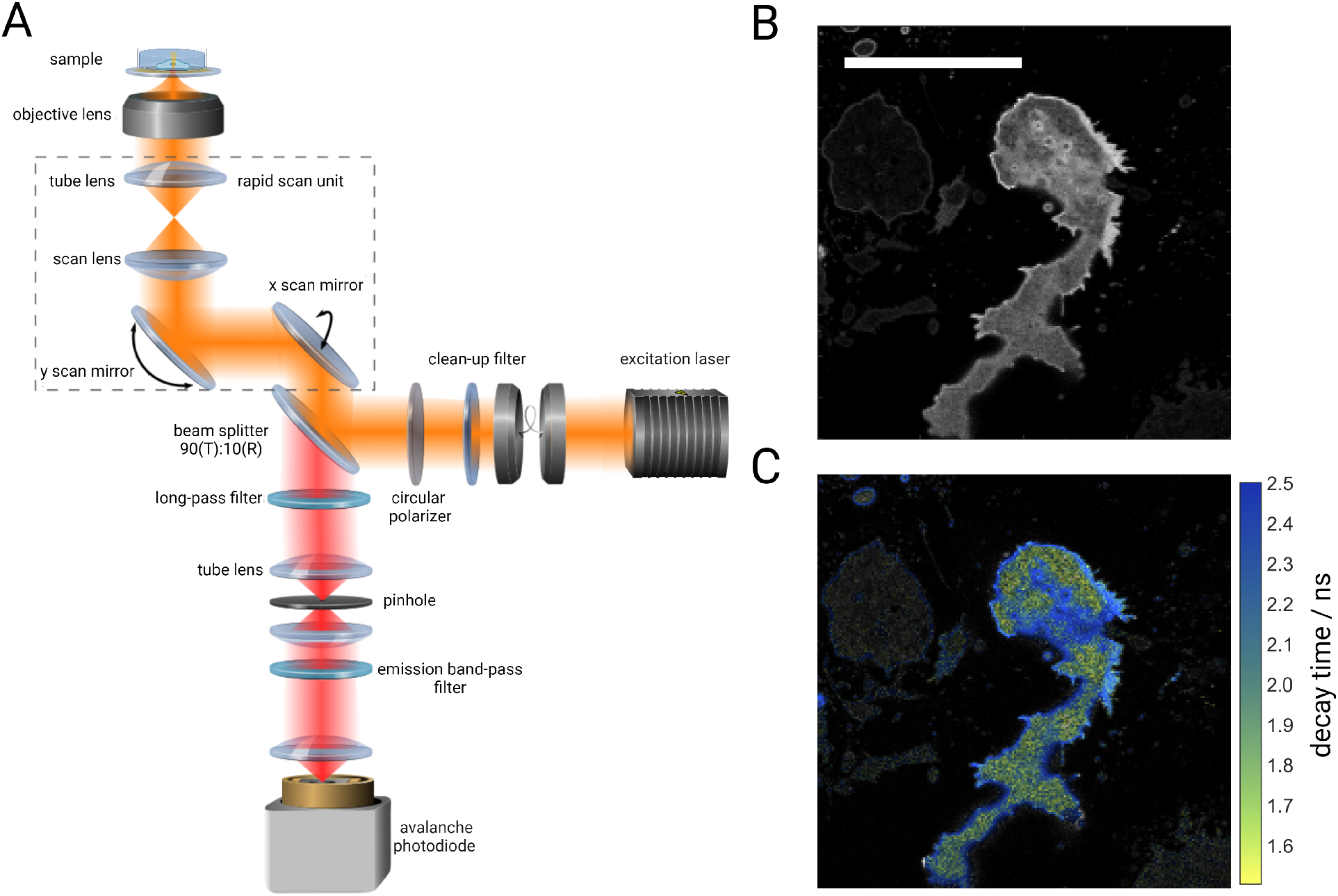
Experimental setup and scheme. **A** demonstrates the confocal fluorescence lifetime imaging (FLIM) microscope equipped with a rapid scanning unit (shown as dotted lines). A pulsed laser source is used for excitation. Excitation beam (shown in orange) is guided through a clean-up filter followed by a beam splitter cube (90:10) which reflects 10% of light to the scan unit. After the scan unit, the beam is focused onto the sample through a high numerical aperture objective lens. Fluorescence light is collected back via the same objective and propagates through the scan unit to the beam splitter cube which transmits 90% of emission and reflects the rest 10%. The transmitted light is then passed through an appropriate long-pass filter and then the beam is focused to a pinhole by a lens. Light after the pinhole is refocused into the active area of a single photon sensitive avalanche photodiode. Optionally, a band-pass filter can be used before the detector to reject residual scattering signal from the metal film in the MIET substrate. **B** and **C** show intensity and fluorescence lifetime images, respectively, of an exemplary *D*.*d*. cell. Scale bar: 10 μm.

We fitted recorded time-correlated single-photon counting (TCSPC) histograms from each pixels to obtain fluorescence lifetime values (see ‘Methods’ for details on lifetime fitting). To convert lifetimes into axial distances, we first measured the (free-space) lifetime of GFP attached to cAR1 for cells on a glass slide without any metal layer. The measured value was *τ*_0_ = 3.1 ns and did not show any systematic variation across cells. Together with the fluorescence quantum yield value of GFP of *ϕ* = 0.79 ^45^, we computed the lifetime-on-distance MIET calibration curve for cells on gold-coated glass cover slides (see ‘Methods’ and supplementary section S3 for MIET curve calculation).

Figure 2A presents reconstructed height images obtained from FLIM images that were converted using the pre-calculated MIET lifetime-to-height calibration curve. Shown are cells in their developed (left panel) and vegetative stage (right panel) (see supplementary section S3 for FLIM images and movies S2, S2e, S3, and S3e). We observe a mean axial distance of 47 ± 8 nm for the cell in its vegetative stage, while the cell in its developed stage approaches the surface much closer, exhibiting a mean axial distance of 27 ± 3 nm (see Figure 2B). The much wider distribution of height values at the vegetative stage indicates significant height fluctuations for this stage. On the contrary, the developed cell pulsed with cAMP shows stronger adherence to the substrate reducing the observed temporal height fluctuations. Furthermore, we observed the formation of pseudopodia in the developed cell and its subsequent chemotactic spreading on the surface (see SI Movie S2) which is, as expected, not observed in the vegetative cell. Another observation is that height values for the vegetative cell are non-uniform across the cell. In particular, they are higher at boundaries as compared to other areas. In contrast, the developed stage exhibits largely homogeneous axial distances throughout a cell. Our experiments confirmed the change of the adhesion of *D*.*d*. cells during the development time and show the difference in terms of axial distances.

**Figure 2:**
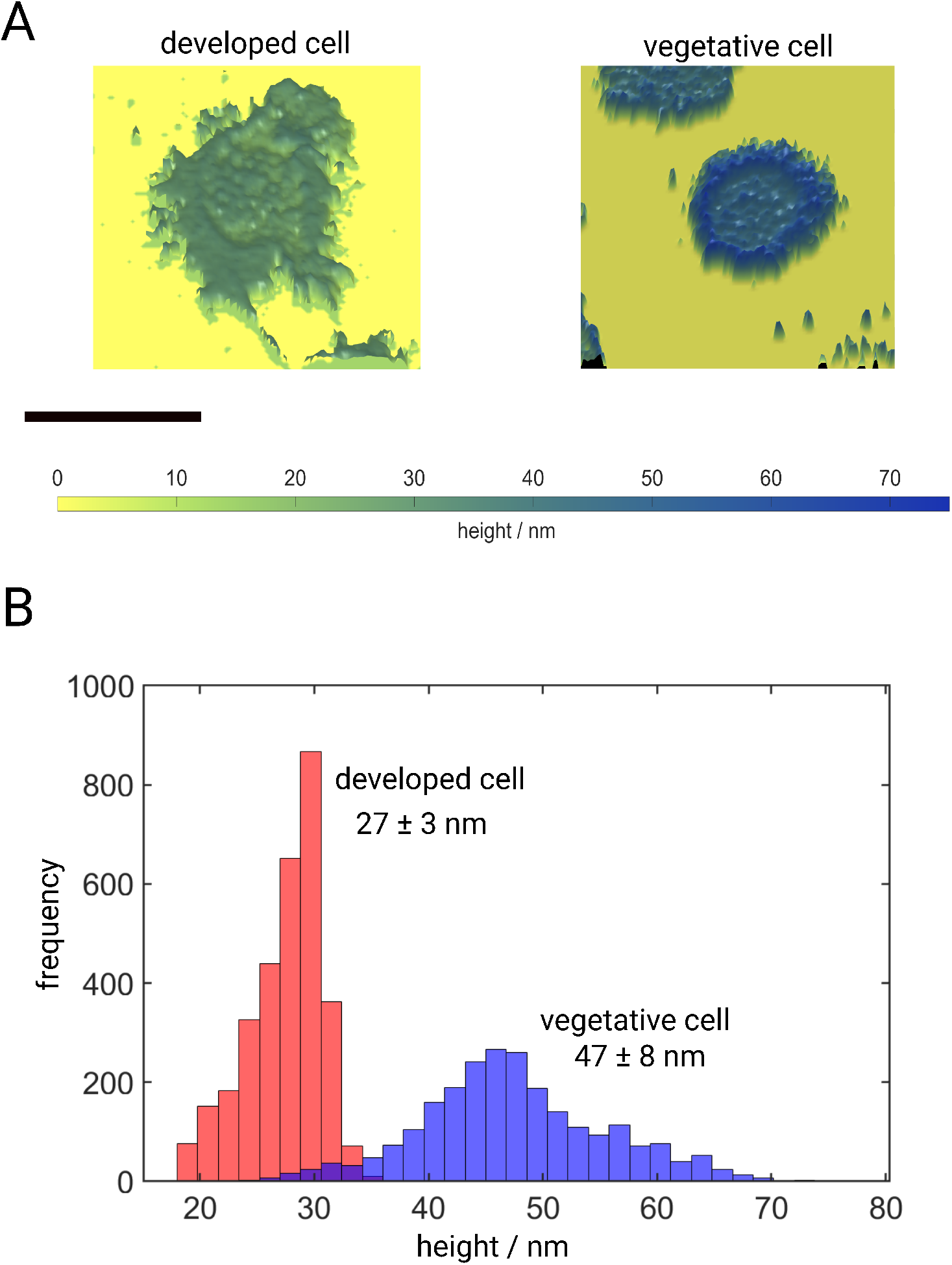
Live-cell MIET imaging of *D*.*d*. cells. **A** presents images of axial distances of a pulsed developed cell (left) and a vegetative cell (right), see also supplementary movies S2 and S3. Scale bar: 10 μm. **B** Histograms visualizing height profiles of the same cells as shown in **A**. The vegetative cell exhibits a mean height of 47 ± 8 nm, which is almost twice the height of the developed cell with 27 ± 3 nm.

FLIM images of *D. d*. cells (see supplementary figure S8) were recorded at a frame rate of 20 Hz (see section *Methods*). However, pertaining to the requirement of a reasonable photon budget for fluorescence lifetime fitting, further analysis was done with a 4x frame binning. Figure 5 A-D illustrates time-lapse images of a representative *D*.*d*. cell in its developed stage (pulsed with cAMP) over a span of 2 s at intervals of 500 ms. False color scale indicates axial distance values. As can be seen in the insets, our live-cell MIET imaging captures fast temporal changes in the morphology of membrane protrusions and their heights.

**Figure 3:**
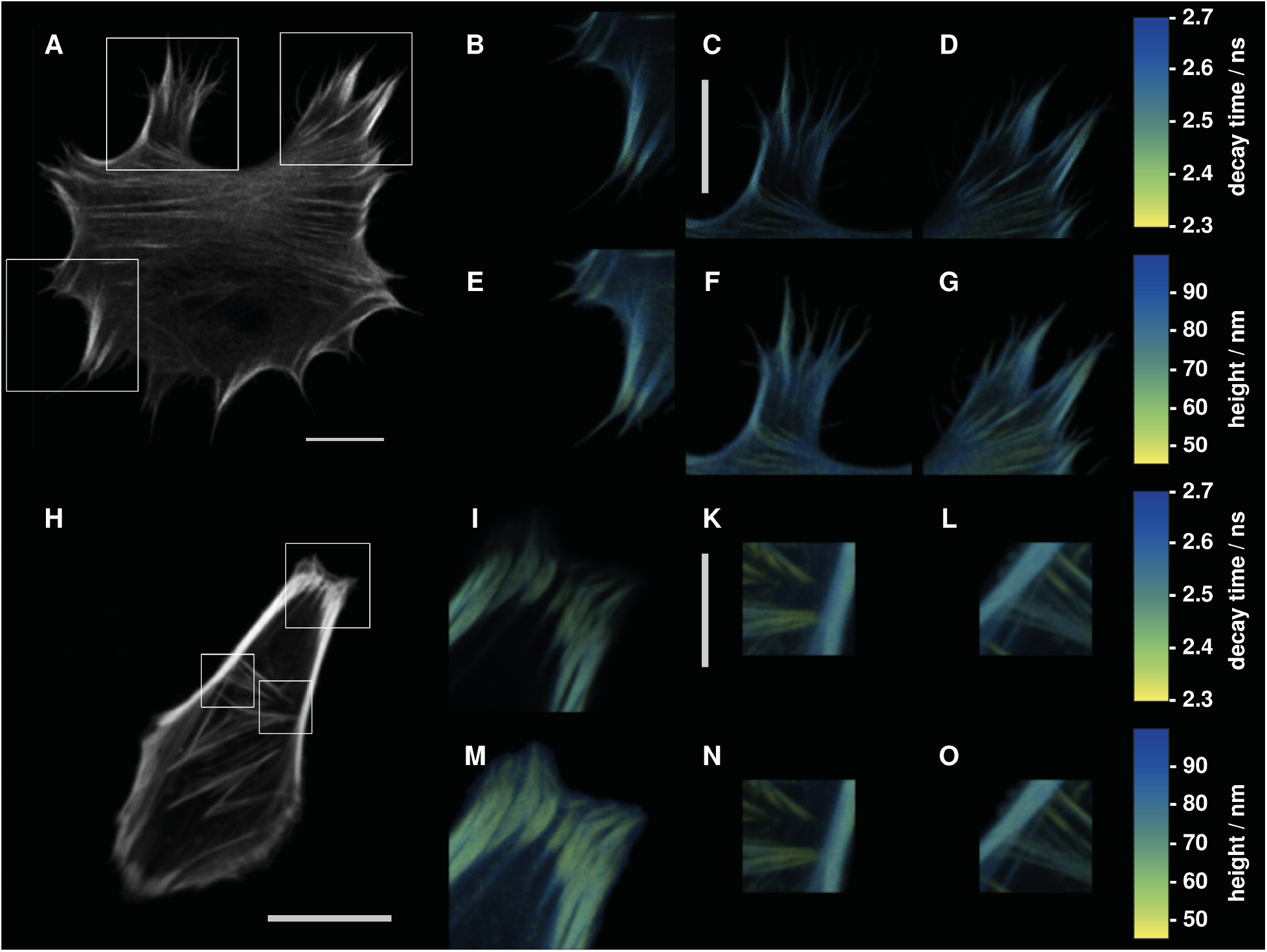
**A**-**G**: hMSC cell 24 h after transfection. **A** shows a fluorescence intensity image of the whole living cell. **B**–**D** show fluorescence lifetime images for different regions of interest (ROIs) shown in **A** (white squares). **E**–**G** are height maps computed from the lifetime images of **B**–**D. H**-**O**: SAOS2 cell 48 h after transfection. **H** show a fluorescence intensity image of the whole living cell. **I**–**L** show fluorescence lifetime images for the ROIs indicated in **H. M**–**O** are height maps computed from the lifetime images of **I**–**L**. All scale bars are 10 μm long.

**Figure 4:**
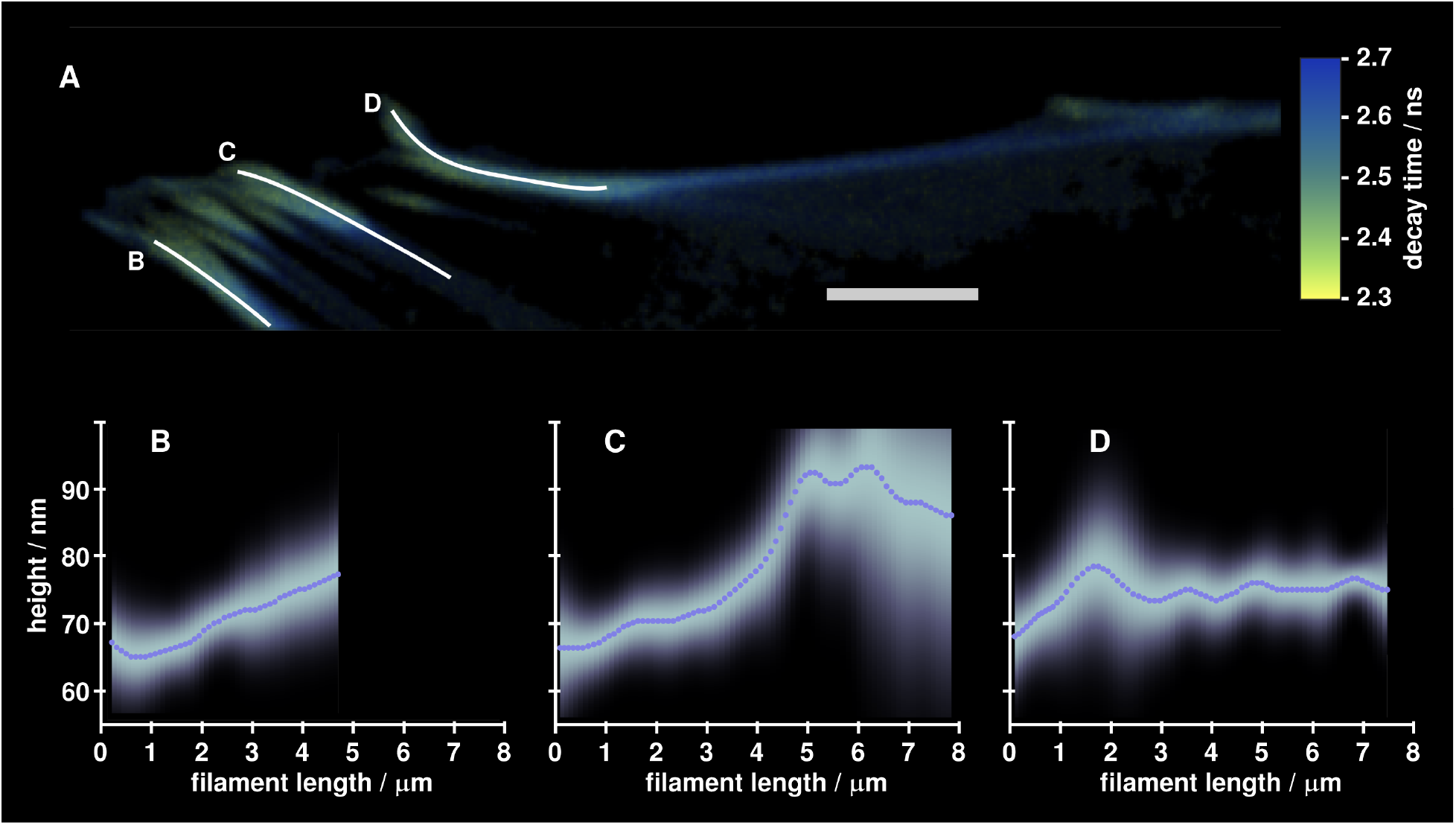
SAOS2 cell 48 h after transfection. **A** shows a fluorescence lifetime image of a cell region showing the presence of stress fibers. Three three annotated fibers were chosen for further analysis. Scale bar is 5 μm. **B**–**D** present show height curves along the three filaments. Dotted lines are mean values, while the density plots around these lines show corresponding error distributions.

**Figure 5:**
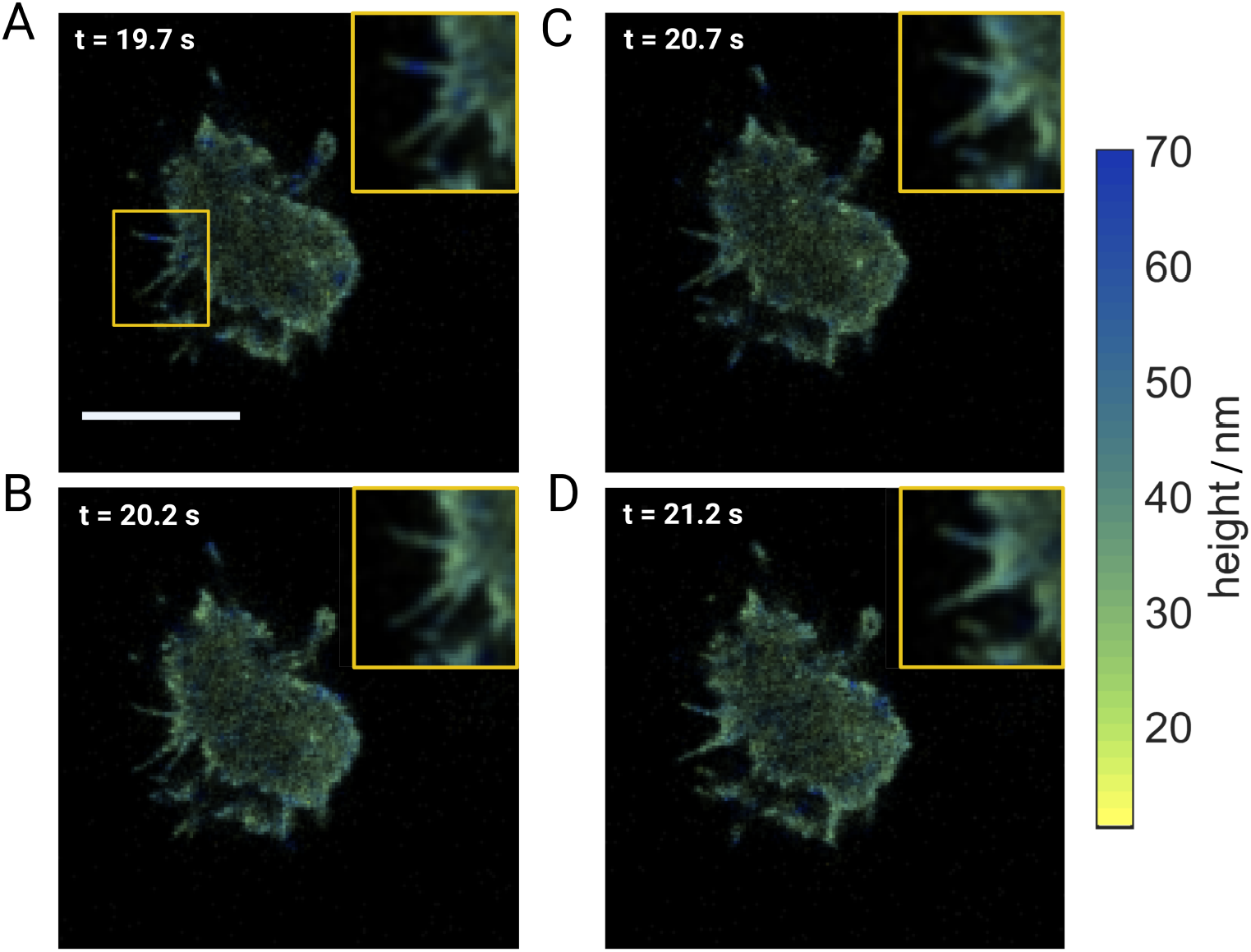
Live cell MIET imaging of *D*.*d*‥ **A - D** Temporal evolution of the morphology of membrane protrusions and axial distances of a *D. d*. cell pulsed with cAMP at its developed stage. Original images were captured at a frame rate of 20 Hz and then binned to 5 Hz. Insets at the top right corners show magnifications of the ROI indicated in the first frame (top left) and detail subtle morphological changes over a time span of 1.5 s (*t* = 19.7 s to *t* = 21.2 s).

The second system studied with FP-based live-cell MIET imaging was the three-dimensional architecture of actin stress fibers in mammalian osteosarcoma cells (SAOS-2) and in human mesenchymal stem cells (hMSCs). The force transmission from contractile acto-moysin stress fibers through focal adhesions to the cells’ surroundings is a topic of intense research^46^. While much is known about structure formation and dynamics of stress fibers in two dimensions^47–49^, the three-dimensional arrangement of proteins in focal adhesions was resolved only recently ^26, 50^. However, these studies were all done exclusively in fixed cells.

We have performed MIET imaging of stress fibers in live SAOS-2 cells 48 hrs after transfection, and Figure 4 shows images of stress fibers emerging from focal adhesions. Brightness in these images corresponds to fluorescence intensity, while color encodes the fluorescence lifetime. Height profiles for the three annotated fibers in panel A are shown in panels B through D. This allows to reconstruct the 3D structure of stress fibers, which is important to fully understand mechanical cell-matrix interactions, in particular the direction of generated and transmitted forces. For the three analyzed stress fibers, actin heights above substrate vary from 60 nm to 100 nm which is in good agreement with fixed-cell measurements using interferometric PALM (iPALM)^50^ and MIET imaging^51^. Our results show very shallow inclination angles of stress fibers with respect to the substrate surface (around 1°), which is in good agreement with our recently reported values using fixed cells and synthetic dyes^26^.

Restructuring of the cytoskeleton and focal adhesions can occur on timescales ranging from minutes to hours. It is thus imperative to observe these cellular processes quantitatively over long periods of time. Using our MIET imaging setup, we were able to image living cells over 7 hours, see Supplemental Video S1 and Supplementary information for details.

## 3 Discussion

Our results demonstrate the feasibility and capabilities of MIET microscopy for axial super-resolution live-cell imaging with fluorescent proteins. We showed this for two different types of mammalian cells (hMSCs and SAOS-2) and for the slime mold *D. discoideum*, using different fluorescent proteins (mScarlet and GFP). We achieved an axial resolution of a few nanometers at an effective image rate of 5 Hz, which is only limited by the brightness and density of the fluorescent proteins but not the MIET imaging microscope itself. Furthermore, we presented long-term measurements of up to 8 hrs, demonstrating the low phototoxicity of MIET imaging, which offers the possibility to follow and quantify cellular processes with extreme axial resolution over long time scales. In particular, this will be of considerable interest for the study of the structure and dynamics of the cytoskeleton and of the focal adhesion machinery.

An important result is the demonstration that MIET imaging works well with FP labeling. FPs are on average less bright than organic dyes, and do often show a non-mono-exponential fluorescence lifetime decay. Nonetheless, our results show that both these limitations still allow for decent live-cell MIET imaging with excellent signal-to-noise ratio and superior axial resolution. Considering different possible origins for the bi-exponential fluorescence decay we could show that we accurately estimate the radiative rate based on the average fluorescence decay time 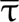 (see SI). For typical FPs, the error of the determined height based on is considerably less than 2 nm. This makes MIET imaging an ideal microscopy modality for a wide range of applications requiring live-cell imaging with fluorescent proteins. When compared to non-fluorescent interferometric techniques such as quantitative phase imaging^52^ that deliver nanometer axial resolution, MIET imaging provides the specificity of fluorescence in addition to a similar axial resolution. Moreover, because MIET imaging is mostly insensitive to refractive index variations, it is not affected by inhomogeneous protein and cell organelle distributions. Taken together, we believe that live-cell MIET imaging with fluorescent proteins can be an important addition to the toolbox of axial super-resolution fluorescence microscopy techniques for cell biology.

## Data Availability

The data that support the plots within this paper and other findings of this study are available from the corresponding authors upon reasonable request.

## Code Availability

All Matlab routines and codes used for data analysis of this study are available from the corresponding authors upon reasonable request.

## Methods

### Cell culture

Human mesenchymal stem cells (hMSCs) (Lonza Ref. #PT-2501, Lot# 603525) were grown in T75 cell culture flasks (Corning Inc., New York, NJ, USA, 430641U) in low glucose DMEM (Gibco, Thermo Fisher Scientific Inc., Waltham, MA, USA, 31885-023) supplemented with 10% fetal bovine serum (Sigma-Aldrich Co., St. Louis, MO, USA, F2442-500ML) and 1% antibiotics (penicillin/streptomycin, Life Technologies, Thermo Fisher Scientific Inc., Waltham, MA, USA, 15140-122) at 37 °C and 5% CO_2_ and passaged every 2-3 days (cells of passage #6 were used in this study).

SAOS-2 cells (DMSZ, ACC 243, RRID:CVCL 0548) were grown in T75 cell culture flasks (Sarstedt, Nümbrecht, Germany, 833.911.002) in McCoy’s 5A (Gibco, Thermo Fisher Scientific Inc., Waltham, MA, USA, 26600-23) supplemented with 15% fetal bovine serum (Sigma-Aldrich Co., St. Louis, MO, USA, F2442-500ML) and 1% antibiotics (penicillin/streptomycin, Life Technologies, Thermo Fisher Scientific Inc., Waltham, MA, USA, 15140-122) at 37 °C and 5% CO_2_ and passaged every 4 days (passage #13 was used in this experiments).

carA-GFP *D*.*d*. cells were cultivated in HL5 medium (Formedium) at 22 °C on polystyrene Petri dishes (Primaria, Falcon, BD Becton Dickinson) and passaged every 2-3 days. As long as nutrients are available (HL5 medium), *Dd* cells proliferate as unicellular amoeba and are defined as vegetative cells. When the cells deplete their food source and start to starve (in our case, the HL5 medium is removed and replaced by buffer), they enter a developmental cycle. For preparation of experiments, cells were starved in shaking phosphate buffer at 150 rpm (PB, 2 g KH_2_PO_4_ and 0.36 g Na_2_HPO_4_ · 2H_2_O per 1 L, pH 6) for 5h at a density of 2 · 10^6^ cells/mL. The shaking culture was pulsed with 50 nM cAMP (Sigma) every 6 min over the course of the starvation time. After the corresponding starvation time the cells were harvested and washed in PB. An aliquot of the cell suspension was applied to the experimental set up and the cells were allowed to spread on the glass substrate for 10 min at room temperature before starting the experiment. For experiments with vegetative *D*.*d*. cells they were detached from the Petri dish bottom, washed twice with PB and applied directly to the experimental set up without any additional starvation time.

### Transfection

Cells were transfected with the plasmid pLifeAct mScarlet-i N1, a gift from Dorus Gadella (Addgene plasmid #85056, RRID:Addgene 85056), using the Lonza 4D Nucleofector (Lonza, Cologne, Germany, AAF-1002B & AAF-1002X). hMSCs were transfected using the P1 buffer (Lonza, Cologne, Ger, V4XP-1012) and pulse code ‘FF-104’. SAOS-2 cells were transfected using the SF buffer (Lonza, Cologne, Ger, V4XC-2012) and pulse code ‘DS-150’. Subsequent to transfection cells were seeded onto prepared dishes at 80.000 cells per well. Dead cells were removed by rinsing with medium 12 h after transfection.

The cell line carA-GFP were derived from the axenically growing strain *D*.*d*. AX3. They were kindly provided by G. Gerisch (Max Planck Institute for Biochemistry, Martinsried, Germany).

### Sample Preparation

For SAOS-2 and hMSC measurements, glass coverlips of 25 mm diameter (VWR, Darmstadt, Ger, ECN631-1584) were covered with 2 nm Ti, 15 nm Au, 1 nm nm Ti, 20 nm nm SiO_2_. Both, gold coated and regular glass coverslips were cleaned and functionalized with collagen-I using a hetero-bifunctional cross-linking chemistry as described earlier^53^. In brief, all slides were plasma cleaned for 15 min, sonicated first using 99% EtOH, second using 2% APTES (Sigma-Aldrich Co., St. Louis, MO, USA, 440140) in EtOH and treated with a 0.5% gluteraldehyde solution (Sigma-Aldrich Co., St. Louis, MO, USA, C7651). Glasses were rinsed twice with PBS and once with HEPES buffer. Sulfo-SANPAH (Thermo Fisher Scientific Inc., Waltham, MA, USA, 22589) was added to the surface and activated under UV for 10 min. A solution of 0.2 mg/mL Collagen I (Rat tail Collagen I, Corning Inc., New York, NJ, USA, 354236) in PBS is added and incubated over night at 4 °C. Glasses are rinsed twice with PBS and glued into bottomless ibidi dishes (ibidi, Gräfeling, Ger, DIO01110) using UV-curable glue (NOA68, Norland products inc., Cranbury, NJ, USA, 6801). Prepared dishes are UV sterilized for 2 hours and rinsed three times with PBS before seeding of cells.

For MIET measurements on *D*.*d*. cells, glass coverslips were first sonicated with 1 M KOH and plasma cleaned for 15 minutes. These cleaned coverslips were then coated with 2 nm titanium layer (for better sticking of gold on glass) followed by evaporation of 15 nm gold, 1 nm titanium and 10 nm SiO_2_. Gold-coated coverslips were then glued into bottomless ibidi dishes (ibidi, Gräfeling, Ger, DIO01110) using UV-curable glue (NOA68, Norland products inc., Cranbury, NJ, USA, 6801). Prepared dishes are UV sterilized for 2 hours and rinsed three times with PBS before seeding of cells.

### Imaging

Cells were imaged in Phenol red free DMEM (Gibco, Thermo Fisher Scientific Inc., Waltham, MA, USA, 11880-028) supplemented with 12,5% fetal bovine serum (Sigma-Aldrich Co., St. Louis, MO, USA, F2442-500ML) and 1% antibiotics (penicillin/streptomycin, Life Technologies, Thermo Fisher Scientific Inc., Waltham, MA, USA, 15140-122) at 37 °C, 5% CO_2_ and 80% humidity using a climatic chamber (ibidi, Gräfeling, Ger, 10918) and gas mixer with humidifier column (ibidi, Gräfeling, Germany, # 11922-DS). carA-GFP *D*.*d*. cells were imaged in PB at room temeparture.

### Setup

Fluorescence lifetime images were acquired on a custom-built confocal setup. The excitation light was generated by an 80 MHz pulsed white-light laser (SuperK Power, Koheras) and the wavelength was selected by an AOTF (SpectraK Dual, Koheras). The beam was coupled into a single-mode fibre (PMC-460Si-3,0-NA012-3APC-150-P and fibre coupler 60SMS-1-4-RGBV-11-47, both Schäfter + Kirchhoff GmbH, Germany) and after the fibre recollimated by an objective (UPlanSApo 10× 0.40 N.A., Olympus). After passing a clean-up filter (F37-563 and F49-488 (for *D*.*d*. measurements), AHF), a 90/10 beam splitter was used to reflect the excitation light into the microscope and separate it from the emission. The reflected beam was directed into a laser scanning system (FLIMbee, PicoQuant) and then into a custom sideport of the microscope (Olympus IX73). The three galvo mirrors in the scanning system are imaged onto the backfocal plane of the objective (UApo N 100× 1.49 N.A. oil, Olympus) with 180mm and 90mm achromatic lenses. The sample could be moved by a manual *xy*-stage (Olympus) and a *z*-piezo stage (Nano-ZL100, Mad-CityLabs). Fluorescence in the sample was collected by the same objective and descanned in the scanning system. The fluorescence light that passed the 90/10 beam splitter was then focused onto a pinhole (100 μm, Thorlabs) with a 180mm achromatic lens. Backscattered laser light was blocked by a long-pass filter (568 LP Edge Basic, Semrock and F76-490 for *D*.*d*. GFP measurements). The light was collimated by a 100mm lens and passed through a bandpass filter (FF01-593/40-25, AHF and F37-521 525/45, AHF for *D*.*d*. GFP measurements) before a lens (*f* = 30mm, Thorlabs) focused the light onto the detector (*τ* -SPAD, PicoQuant). The signal of the photon detector was recorded by a TCSPC system (HydraHarp 400, PicoQuant) together with the trigger signal by the laser. FLIM images were recorded with the SymPhoTime 64 software (Picoquant) which controlled the TCSPC system and the laser scanner. Typically, a pixel size of 100 nm was chosen with a pixel dwell time of 20 μs and a TCSPC resolution of 16 ps. For *D*.*d*. GFP measurements, a pixel size of 150 nm was chosen with pixel dwell time of 2 *μ*s and a TCSPC resolution of 16 ps. To monitor the axial focus position, back-reflected light was coupled out from the excitation beam with an additional 90/10 beam splitter and focused (*f* = 200mm, Thorlabs) onto a camera (Guppy GF036B ASG, Allied Vision Technologies).

### Data Analysis

Data analysis was carried out using custom routines written in MatLab (MathWorks). First, scan lines were aligned using a linear shift. Then, TCSPC histograms of individual pixels were generated and corrected for dead-time artefacts.^54^ A mono-exponential fit including a measured instrument response function (IRF) was used to estimate the fluorescence lifetime. The TCSPC curve was fitted with the function

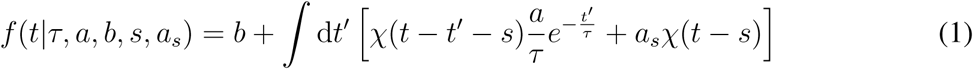

where *χ*(*t*) denotes the instrumental response function of the TCSCP system, *τ* denotes the fluorescence lifetime, *a* denotes the fit amplitude, *b* the background, *a*_*s*_ the scattering amplitude, and *s* the color shift of the IRF. The scattering amplitude accounts for the luminescence of the gold, and the shift *s* was introduced as the IRF depends on the count rate. We found for our detectors that linear shifts was sufficient to account for this effect. The lifetime was converted into a height using the appropriate MIET curve for the sample. Images and movies were created with the lifetime or height information in color, using the intensity (re-scaled to 0−1) as the transparency of the image.

### MIET curve calculation

The theoretical curve for converting measured lifetimes into heights were computed based on the model by Chance, Prock, and Silbey^25, 55, 56^. For the substrate, we assumed a layered system of (from bottom to top): glass, 2 nm titanium, 15 nm gold, 1 nm titanium, and 20 nm silica. The medium of the fluorophores and above was assumed to have the refractive index of water, refractive indices of the metals were taken from literature ^57^. The theoretical MIET curve for GFP in connection to *D*.*d*. measurements were calculated similarly where we assumed a layered system (from bottom to top) glass, 2 nm titanium, 15 nm gold, 1 nm titanium, and 10 nm silica. Please see supplementary information section S3 for theoretical MIET curve.

### Impact of multi-exponential fluorescence decays

Like other methods that take advantage of fluorescence quenching (Förster resonance energy transfer, Stern-Vollmer quenching) the analysis is based on the fluorescence decay time, being the inverse of the rates depopulating the excited state. These are the emission rate *k*_fl_ and non-radiative processes that are summarized into *k*_nr_. The quenching adds a further rate to the total and the analysis is all about quantifying this additional contribution. For this, *k*_fl_ and *k*_nr_ are assumed to be constant and can be obtained from the fluorescence decay time and the quantum yield of the dye in ‘free-space’ conditions. All these processes lead to a strictly mono-exponential decay of the decay allowing seamless analysis of the data. In reality however, one usually observes fluorescence decays that deviate from the expected behavior. Most common are bi-exponential decays. These are usually explained as two populations of slightly different forms of the fluorophore (e.g. in a different local environment, charges near the chromophoric part of the molecule etc.).

Accurate determination of two lifetimes and their respective contribution to the decay is surprisingly hard and requires collecting many photons. Therefore, we estimate the amount of quenching based on the average fluorescence decay time, which can be determined quite simply and reliably. But how do the results of this simplified analysis differ from the true heights? We simulated the biexponential decays based on the rate constants of lifeact-mScarlet and the calculated MIET curve. The average lifetime of these quenched decays were determined and used to calculate the height, again using the MIET curve. The results clearly prove that the analysis based on the average fluorescence decay time adds no significant error to the determined heights of the fluorophore. (see section S.1 in the supporting information)

### Protein expressions for QY measurement

#### lifeact-mScarlet from SAOS-2

For quantum yield measurements, SAOS-2 cells (passage #13) were cultured and transfected as previously mentioned. The fusion protein (Lifeact-mScarlet) was extracted using a RFP-Trap (Chromotek, Planegg – Martinsried, Germany, rta-20). Cells were harvested, lysed, and protein was extracted as advised by vendor. For elution, we bound proteins by adding 50 μl 0.2 M glycine (pH 2.5) under 30 sec of constant mixing. After 2 min centrifugation at 2.500 g supernatant was transferred to a new tube and pH neutralized by addition of 5 μL 1 M Tris base (pH 10.4).

#### lifeact-mScarlet-6His in E. Coli

The lifeact-mScarlet sequence was cut out using NdeI and XhoI sites and inserted into pET24b+ vector (Sigma-Aldrich Co., St. Louis, MO, USA, 39750), yielding a 6-His Tag on the mScarlet. Construct was electroporated into E.coli (BL21-Gold Competend Cells) (Agilent Technologies, Santa Clara, CA, USA, #230130) and plated on LB plates (SigmaAldrich Co., St. Louis, MO, USA, 52062) with kanamycin (Sigma-Aldrich Co., St. Louis, MO, USA, 70560-51-9). Growing cultures were sequenced (Sanger Sequencing, Microsynth Seqlab, Göttingen, Ger) and expressed in 500 ml LB Medium (Sigma-Aldrich Co., St. Louis, MO, USA, 51208) for 5 hours before harvesting. Cultures were spun down and pallets subjected to lysis buffer (50 mM Tris/Cl pH 8.0, 250 mM NaCl, 10 mM *β*-Mercaptoethanol, 1 mM PMSF). LifeactmScarlet-6His was bound to Protino NiNTA Agarose (Macherey-Nagel, Düren, Ger, 745400.25) and washed (50 mM Tris/Cl pH 8.0, 250 mM NaCl, 10 mM *β*-Mercaptoethanol, 20 mM Imidazol). NiNTA beads were given into a empty Protino column (Macherey-Nagel, Düren, Ger, 745400.10) and washed again. Final elution was done with 5 times 1 ml elution buffer (50 mM Tris/Cl pH 8.0, 250 mM NaCl, 10 mM *β*-Mercaptoethanol, 250 mM Imidazol) for 30 min each and samples frozen at −80°C. Samples were checked with a SDS-Page gel.

## Acknowledgements

We are grateful to the Deutsche Forschungsgemeinschaft (DFG, German Research Foundation) for financial support through projects A06 of the SFB 860 (JE,AG,NK) and project B08 of the SFB 755 (FR). JE is grateful to the European Research Council (ERC) via project “smMIET” (grant agreement no. 884488) under the European Union’s Horizon 2020 research and innovation program, and for financial support through Germany’s Excellence Strategy–EXC 2067/1-390729940. We thank the Leibniz Association for financial support through project K76/2017. item[Competing Interests] The authors declare that they have no competing financial interests.

## Correspondence

Correspondence and requests for materials should be addressed to FR and JE.

## Contributions

F. Rehfeldt and J. Enderlein co-designed and co-directed the project. S. Isbaner performed the MIET imaging measurements and data analysis for the SAOS cells and hMSCs. L. Hauke optimized and prepared samples of SAOS cells and hMSCs and assisted the MIET imaging of them. L. Turco and I. Guido provided the carA-GFP *D*.*d*. cells, prepared the samples and assisted the MIET imaging of them. A. Ghosh and N. Karedla performed the MIET imaging measurements and data analysis for the *D*.*d*. cells. I. Gregor and A.I. Chizhik helped with building and maintaining the MIET imaging measurement set-up. A.I. Chizhik performed the nanocavity-based quantum yield measurements of mScarlet. S. Isbaner, L. Hauke, A. Ghosh, F. Rehfeldt and J. Enderlein wrote the main manuscript. All authors were involved with editing and improving the main manuscript and with writing the Supplementary Information.

