## Supplementary material for "Metal-Induced Energy Transfer (MIET) for Live-Cell Imaging with Fluorescent Proteins": SI.pdf

[†]*Equal contribution first authors.*

### S.1. Determining the Molecule's Height Using the Average Fluorescence Decay-time and MIET

To evaluate, whether it is justified to use the average fluorescence decay-time for a simplified analysis of bi-exponential fluorescence decays we consider two scenarios. In the first case, we assume that the two decay rates are caused by different non-radiative rates $k_{\mathrm{nr},1}$ and $k_{\mathrm{nr},2}$ while the fluorescence rate $k_{\mathrm{fl}}$ is the same for the two populations. In the second scenario, we assume that the quantum yield of both populations are the same. Also, in what follows we always assume that the emission spectrum is the same for both populations. The assumed numbers for the simulated results correspond to the range found in the experiments shown in this manuscript. However, the considered distributions are generally wider than found in the real experiments.

**S.1.1. Constant Fluorescence Rate** From these assumption it follows that the decay-times for the two are

$$\tau_1 = {}^1\!/\!_{(k_{\mathrm{fl}}+k_{\mathrm{nr},1})} \text{ , and } \tau_2 = {}^1\!/\!_{(k_{\mathrm{fl}}+k_{\mathrm{nr},2})} \ .$$

The quantum yields are $\eta_1 = k_{\mathrm{fl}}\,\tau_1$ and $\eta_2 = k_{\mathrm{fl}}\,\tau_2$, respectively. If we have mixture with the relative amounts $N_1$ and $N_2$ of both species, we will observe a fluorescence decay as

$$I(t) = A_1 \exp\left(-{}^t\!/\!_{\tau_1}\right) + A_2 \exp\left(-{}^t\!/\!_{\tau_2}\right) \ ,$$

where $A_1 = N_1 * \eta_1$ and $A_2 = N_2 * \eta_2$. The average decay-time and quantum yield of this mixture will be

$$\overline{\tau} = A_1\tau_1 + A_2\tau_2 \text{ , and}$$

$$\overline{\eta} = A_1\eta_1 + A_2\eta_2 \ .$$

These values allow us to calculate the 'MIET curves' for population 1, population 2, and the average of the two. Figure 1 shows the three curves for parameters that are much more pronounced than what is observed in the case of mScarlet.

These curves show how the decay-times drop near the surface, but it is important to note that this is due to an increase of the radiative rate. Therefore, we need to determine the rate $k_{\mathrm{fl}}(z)$ and accordingly the resulting quantum yield $\eta_{1,2}(z)$ as a function of the height. These dependencies are shown in figure 2.

Using the $z$ dependent values of the quantum yields, we can now determine the average decay-

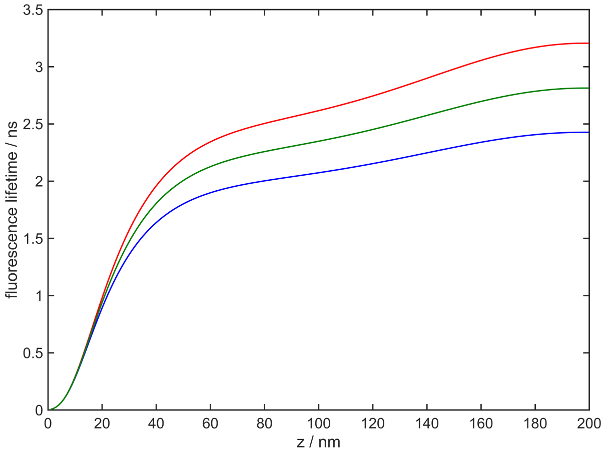


S1.png

Figure 1: Fluorescence decay-time as a function of the height over a MIET surface for different quantum yields. Red: $\tau_1 = 3.00\,\text{ns}$, $\eta_1 = 0.70$, blue: $\tau_2 = 2.30\,\text{ns}$, $\eta_2 = 0.54$, and green: $\overline{\tau} = 2.65\,\text{ns}$, $\overline{\eta} = 0.62$. The green curve was obtained by a mixture of $N_1 = 43.5$ % and $N_2 = 56.5$ %.

time $\overline{\tau}(z)$ that a population of $N_1$ and $N_2$ molecules at the height $z$ will show. Finally, we use $\overline{\tau}(z)$ to compute the height value $\overline{z}$ based on the MIET curve that was calculated from the average free space lifetime (green curve in figure 1). The difference of $\overline{z}$ to the true value $z$ will show the expected error. As shown in the left panel of figure 5 the maximal error is about $\Delta z = 2\,\text{nm}$.

**S.1.2. Constant Quantum Yield** Again, we are starting our analysis considering the decay-times of the two populations

$$\tau_1 = {}^{1}\!/_{(k_{\text{fl},1}+k_{\text{nr},1})} \text{ , and } \tau_2 = {}^{1}\!/_{(k_{\text{fl},2}+k_{\text{nr},2})} \ .$$

The quantum yield is $\eta = k_{\text{fl},1}\,\tau_1 = k_{\text{fl},2}\,\tau_2$, respectively. If we have mixture with the relative amounts $N_1$ and $N_2$ of both species, we will observe a fluorescence decay as

$$I(t) = A_1 \exp\left(-{}^{t}\!/_{\tau_1}\right) + A_2 \exp\left(-{}^{t}\!/_{\tau_2}\right) \ ,$$

where $A_1 = N_1$ and $A_2 = N_2$, and the average decay-time being $\overline{\tau} = A_1\tau_1 + A_2\tau_2$.

Figure 3 shows the 'MIET curves' of population 1, population 2, and the average of the two using parameters that are much more pronounced than what is observed in the case of mScarlet. Next, we determine the rates $k_{\text{fl},1}(z)$ and $k_{\text{fl},2}(z)$ as well as the resulting quantum yield $\eta_{1,2}(z)$ as a

S2a.png S2b.png

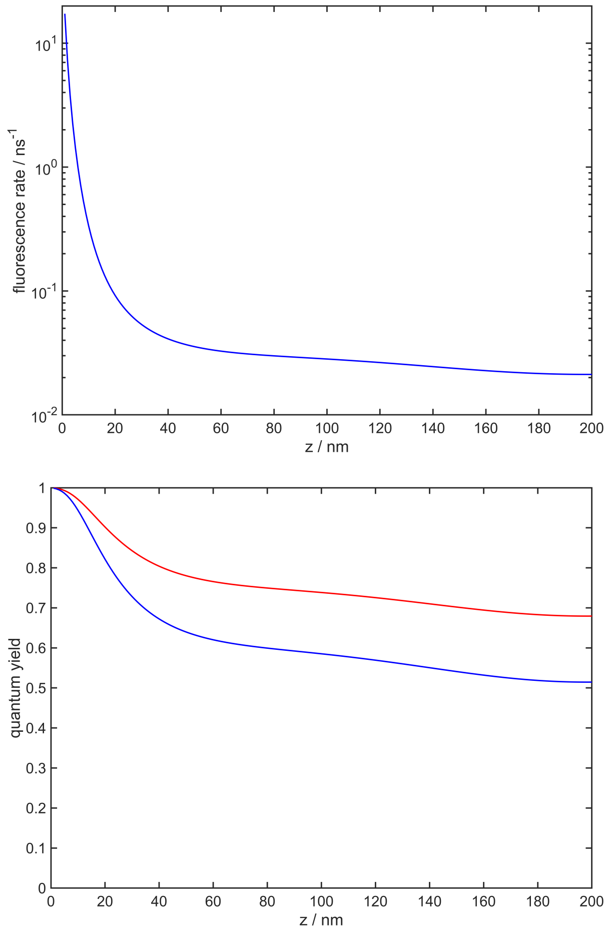


Figure 2: Emission rate $k_{\text{fl}}(z)$ (left) and fluorescence quantum yield $\eta(z)$ (right) as a function of the height over a MIET surface. Red: $\tau_1 = 3.00\,\text{ns}$, $\eta_1 = 0.70$, blue: $\tau_2 = 2.30\,\text{ns}$, $\eta_2 = 0.54$. The given numbers hold for the molecules in homogeneous solution. Because for both molecules, $k_{\text{fl}}$ is the same in the solution, this is also true above the surface.

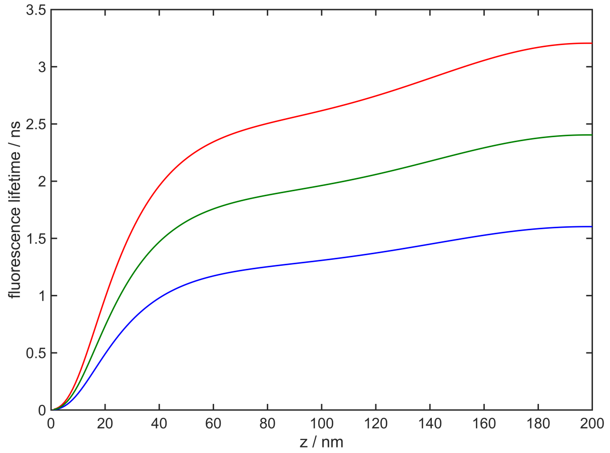


S3.png

Figure 3: Fluorescence decay-time as a function of the height over a MIET surface for the following parameters: $\eta = 0.70$, $\tau_1 = 3.00\,\mathrm{ns}$ (red), $\tau_2 = 1.50\,\mathrm{ns}$ (blue), and $\overline{\tau} = 2.25\,\mathrm{ns}$ (green).

function of the height. These dependencies are shown in figure 4. Because for both molecules the quantum yield is the same in the solution, this is also true above the surface. Using the $z$ dependent values of the quantum yields, we can now determine the average decay-time $\overline{\tau}(z)$ that a population of $N_1$ and $N_2$ molecules at the height $z$ will show. Finally, we use $\overline{\tau}(z)$ to compute the height value $\overline{z}$ based on the MIET curve that was calculated from the average free space lifetime (figure 3). The difference of $\overline{z}$ to the true value $z$ will show the expected error. As shown in the right panel of figure 5 there is no error in the determination of the height. We therefore can safely assume that the average lifetime introduces no significant error in the MIET evaluation.

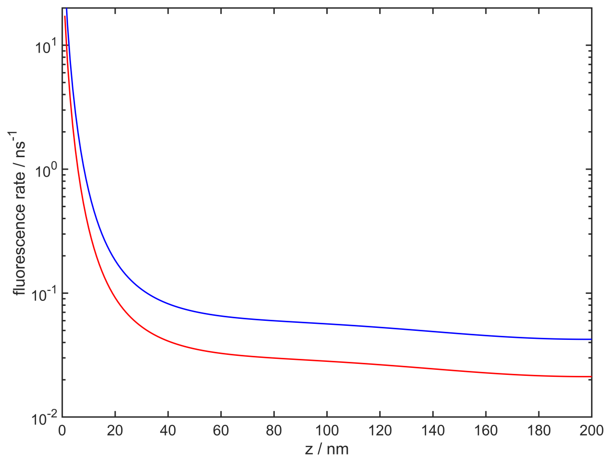


S4a.png S4b.png

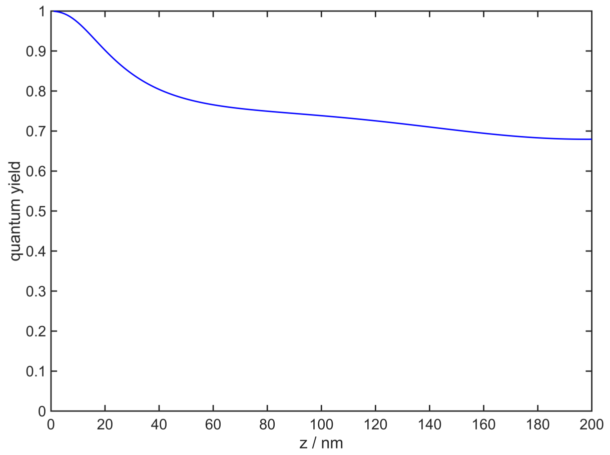


Figure 4: Emission rate $k_{\mathrm{fl}}(z)$ (left) and fluorescence quantum yield $\eta(z)$ (right) as a function of the height over a MIET surface. Free space quantum yield is $\eta = 0.70$. Red: $\tau_1 = 3.00\,\mathrm{ns}$, blue: $\tau_2 = 1.50\,\mathrm{ns}$. The given numbers hold for the molecules in homogeneous solution.

S5a.png S5b.png

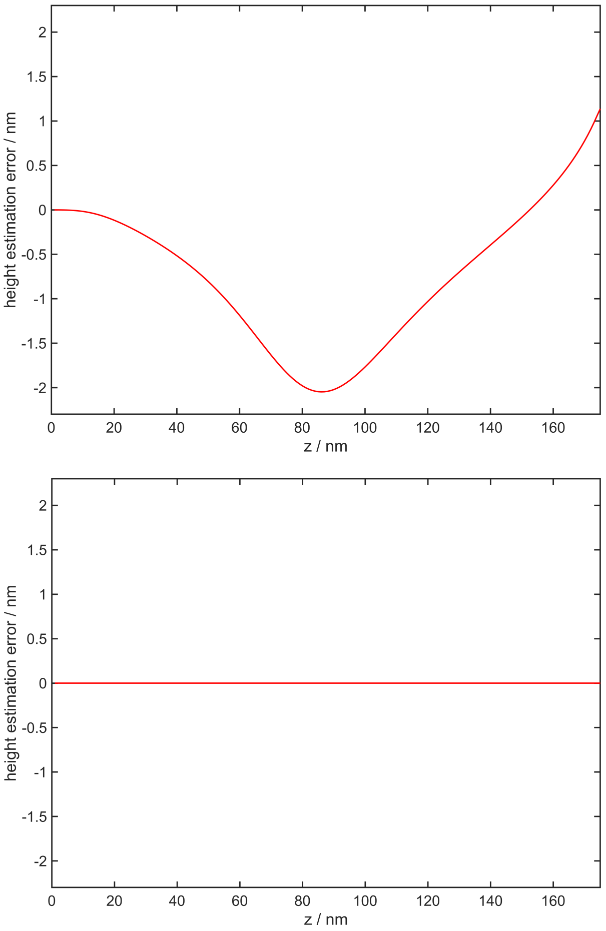


Figure 5: Error of the estimated height ($\Delta z = \overline{z} - z$) due to analysis based on the average fluorescence decay-time. Left: assuming same fluorescence rate; right: assuming same fluorescence quantum yield of both populations.

### S.2. mScarlet lifetime comparison

For some experiments it may be difficult or non-desirable to transfect the cells with a suitable fluorescent protein. In these cases, a viable option might be to micro-inject the protein into the cells. In order to assess this possibility, we compared the fluorescence lifetimes of lifeact-mScarlet in SAOS-2 cells (passage #16, ACC 243, DSMZ). One batch of cells was transfected as described earlier and FLIM images were recorded 24 h post transfection. A second batch was grown in

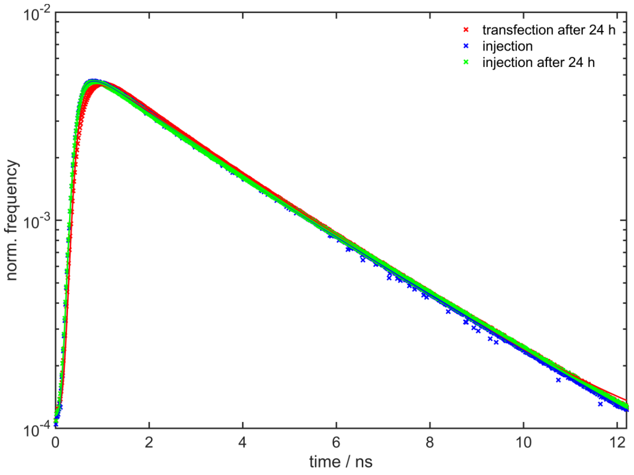


S6a.png

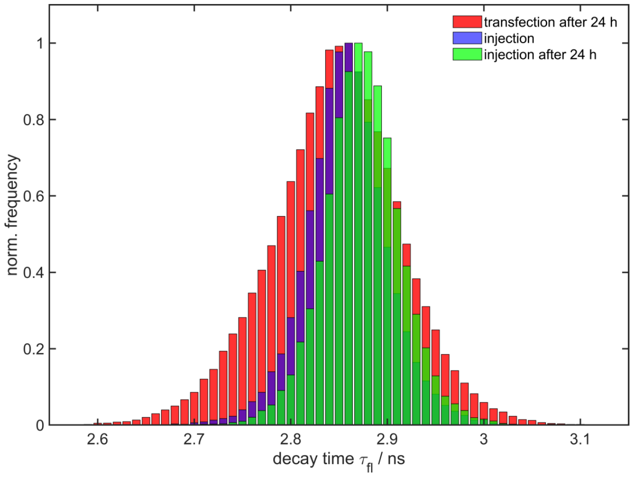


S6b.png

Figure 6: Left: Fluorescence decays of lifeact-mScarlet in hMSC cells. The fitted average decay-times are 2.68 ns for the transfected cells, 2.81 ns for the cells directly after micro-injection of purified protein, and 2.86 ns at 24 h after micro-injection. Right: Histograms of pixel-integrated fluorescence decay-times of the same samples. The histograms show mean decay-times of 2.85 $\pm$ 0.07 ns, 2.86 $\pm$ 0.04 ns, and 2.87 $\pm$ 0.04 ns, respectively. One may note, that the distribution in the case of the transfected cells is broader and shows a tail towards lower decay-times as compared to the micro-injected cells.

parallel without transfection. Prior to the measurement we used a Transjector 5246 (Eppendorf, Hamburg, Ger, 5246 01084) and Femtotips (Eppendorf, Hamburg, Ger, 930000035) to injected about 500 pL of a purified solution of lifeact-mScarlet-$His_6$ (0.78 mg/ml in elution buffer (50 mM Tris/Cl pH 8.0, 250 mM NaCl, 10 mM $\beta$-Mercaptoethanol, 250 mM Imidazol) with 2% sucrose) into a number of cells. FLIM images of these cells were recorded in the same way as for the transfected cells. One set of images was recorded directly after micro-injection, a second set was recorded 24 h after micro-injection.

A comparison of the obtained results is shown in figure 6. The obtained fluorescence decay times agree very well, showing that both options to stain the living cells are work. The distribution of the decay-times in the transfected cells is broader than for the micro-injected cells. Reasons for this might be the more uniform concentration of the protein in the micro-injected cells, or a more homogeneous state of maturation of the purified fluorescent protein. In the transfected cells one has a continuous synthesis (and degradation) of proteins in the cells. This may increase the inhomogeneity of the fluorescence properties. In this way, micro-injection could be advantageous compared to transfection.

### S.3. MIET curve calibration for *D.d* measurements

S7.png

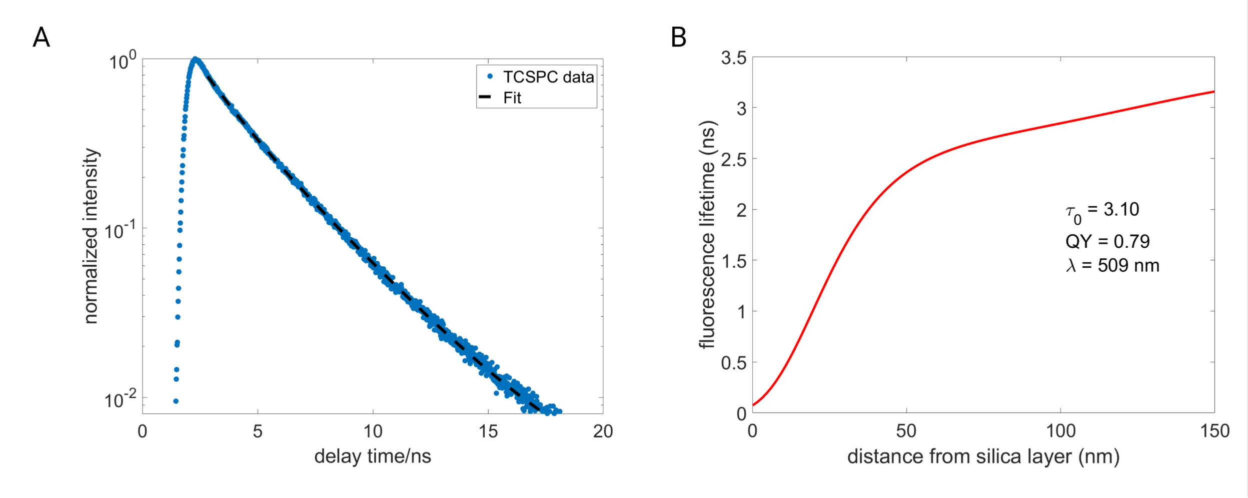


Figure 7: **A.** Free space lifetime of GFP in *D.d.* cells. TCSPC histogram on glass and tail-fit of the decay is shown. We performed a bi-exponential tail-fit to obtain lifetime values of 3.3 ns and 1.3 ns with fit amplitudes of 0.88 and 0.12 respectively. The average lifetime was calculated to be 3.1 ns which was used to compute the MIET calibration curve shown in **B**. **B** MIET calibration curve as obtained from experimentally determined free-space lifetime ($\tau_0$) = 3.1 ns, quantum yield (QY) = 0.79 and at emission wavelength ($\lambda$ = 509 nm).

**S.3.1. Fluorescence lifetime values of GFP in *D.d* measurements** Additional FLIM images and corresponding lifetime histograms for GFP-labeled *D.d* are presented in Fig. 8.

### S.4. Additional data and videos

**S.4.1. SAOS video** Video S1 shows part of a migrating SAOS cell 96 hours post transfection with lifeact-mScarlet. Continuous liveMIET measurement was done with 600 x 600 pixel (corresponding to 120 µm × 120 µm) field of view and stopped after 7 hours.

S8a.png

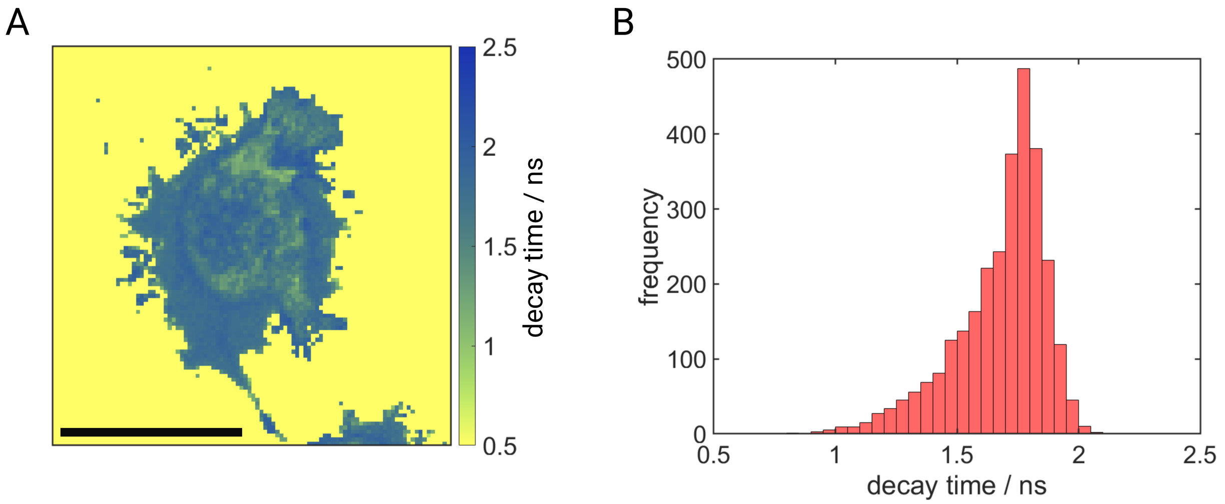


S8b.png

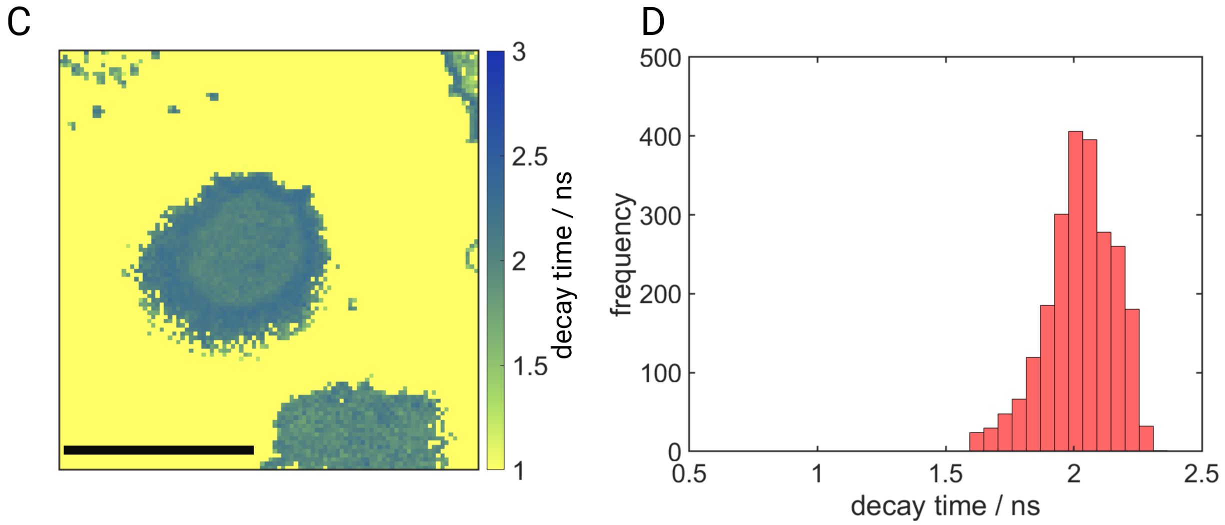


Figure 8: **A.** FLIM image from of a recorded spreading *D.d.* cell in its developed stage pulsed with cAMP. Axial distances of the same cell are presented in Figure 2 in the main text. **B**. Fluorescence lifetime values as obtained from **A**. **C.** FLIM image from a *D.d.* cell in its vegetative stage. Axial distances of the same cell are presented in Figure 2 in the main text. **D**. Fluorescence lifetime values as obtained from **C**. Scale bar 10 $\mu$m

**S.4.2. *D. discoideum* videos** Videos S2 and S2e analysis show proliferation and membrane dynamics of developed *D.d.* cells pulsed with cAMP. Movie S2 corresponds to Figure 2 displayed in the main text. Video S2e illustrates the error values in height determination per pixel of movie S2. False color scale represents axial distances from the gold surface. Videos S3 and S3e illustrate dynamics of *D.d.* cells in its vegetative stage. Video S3 corresponds to Figure 2 displayed in the main text. Video S3e illustrates the error values in height determination per pixel of movie S2. False color scale represents axial distances from the gold surface. All Videos are played at 30 fps for better visualization.

**S.4.3. TCSPC data of other fluorescent proteins** Figures 9 – 11 show TCSPC histograms of other fluorescent proteins that were assessed for our studies on live-cell MIET.

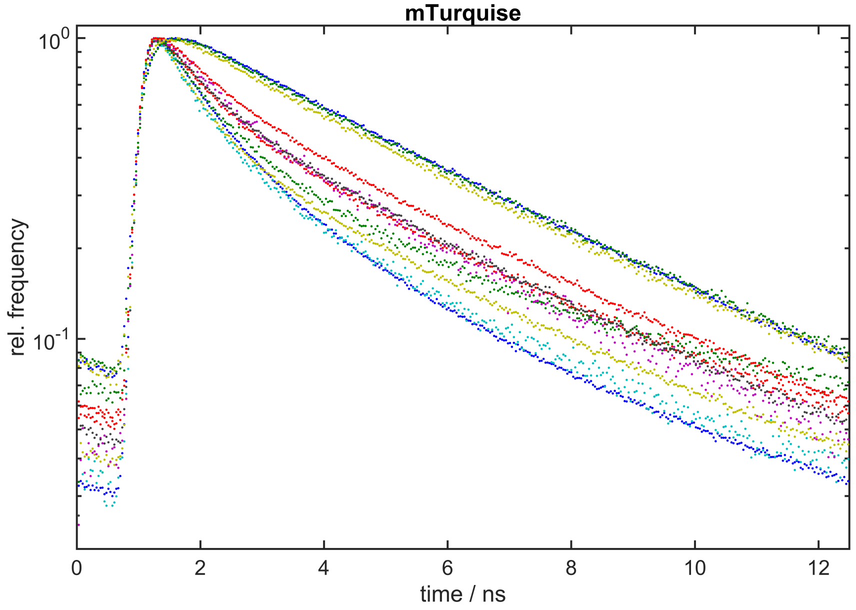


S9.png

Figure 9: Fluorescence decays of lifeact-mTurquise in hMSC cells. Some cells show a mono-exponential decay with a decay-time of about 4.2 ns. Many other cells show bi-exponential decays with a faster component of about 0.7 ns and a slower component of about 3.5 ns. The respective amplitudes vary from cell to cell.

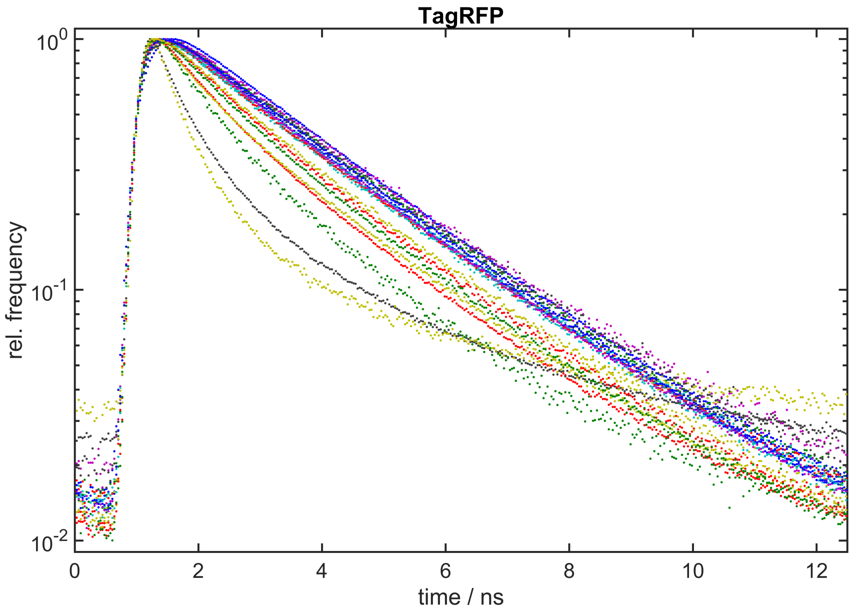


S10.png

Figure 10: Fluorescence decays of lifeact-TagRFP in hMSC cells. Some cells show a mono-exponential decay with a decay-time of about 2.4 ns. Many other cells show bi-exponential decays with a faster component varying between 1.5 ns and 0.7 ns.

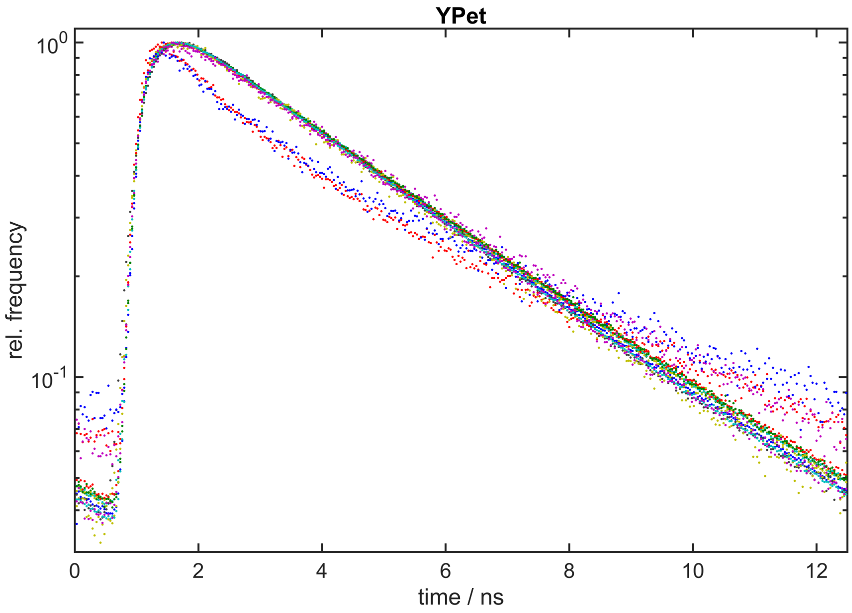


S11.png

Figure 11: Fluorescence decays of lifeact-YPet in hMSC cells. Most cells show a mono-exponential decay with a decay-time of about 3.5 ns. Some cells show bi-exponential decays with a faster component of about 0.8 ns and a slightly prolonged slower component of about 3.8 ns.
